# Self-paced treadmills do not allow for valid observation of linear and non-linear gait variability outcomes in patients with Parkinson’s disease

**DOI:** 10.1101/2020.03.16.993899

**Authors:** Maryam Rohafza, Rahul Soangra, Jo Armour Smith, Niklas König Ignasiak

## Abstract

**Background:** Due to the imposed constant belt speed, motorized treadmills are known to change linear and non-linear gait variability outcomes. This is particularly true of patients with Parkinson’s disease where the treadmill can act as an external pacemaker. Therefore, the use of treadmills is generally not recommended when quantifying gait variability. Self-paced treadmills allow for updating the belt speed relative to the walking speed of the subject and might, therefore, be a useful tool for the collection of long consecutive walking trials, necessary for gait variability observations.

**Research question:** To validate gait variability measures collected on a self-paced treadmill as compared to overground walking.

**Methods:** Thirteen healthy subjects and thirteen patients with Parkinson’s disease performed 5 – 8 minute long walking trials: overground, on a treadmill at a constant speed, as well as in three different self-paced treadmill modes. Stride times and stride lengths were recorded using a validated IMU-system and variability was quantified using the coefficient of variation, sample entropy, and detrended fluctuation analysis. Overground and treadmill trials were compared using Pearson’s correlation coefficient, method error, and Bland and Altman analysis.

**Results:** For healthy subjects, the self-paced treadmill resulted in increased correlation coefficients of 0.57 – 0.74 as compared to a constant speed treadmill. Correlation coefficients for stride length variability between overground and treadmill walking were not significant. For patients, generally, large errors of 33-40% of stride time variability were observed between overground and treadmill walking. Stride length variability is most similar at a constant belt speed and shows errors of 14-39%.

**Significance:** Despite an improvement of temporal gait variability validity in the self-paced mode for healthy subjects, the large systematic and random errors between overground and self-paced treadmill walking prohibit meaningful gait variability observations in patients with Parkinson’s disease using self-paced treadmills.

## Introduction

Walking is one of the most common activities of daily living and essential for independent and self-determined life [1-3]. With increasing age as well as under the influence of disease, walking deteriorates, and unsafe walking patterns have been associated with the occurrence of falls in the elderly and diseased populations [4, 5]. Therefore, the assessment of gait patterns has become an important clinical outcome. In particular, measures to quantify variability are in the focus, as they allow to distinguish between diseased and non-diseased populations [6, 7], can evaluate treatment success [8] and might enable prediction of future adverse events such as falling [9-11].

Generally, gait variability can be quantified by the overall magnitude of spatial and temporal variability in the data [12-14]. This is commonly done by estimating inter-cycle variations in gait derived outcomes, for example, the standard deviation of multiple stride lengths or stride times. However, this linear statistical approach incorrectly assumes that strides are independent of each other [15, 16]. In order to overcome this limitation, non-linear methods have been developed that account for the interdependency of consecutive strides, and those measures, therefore, more accurately quantify the temporal structure of gait variability [15, 17-19]. Two commonly used non-linear methods to evaluate the regularity and self-similarity in discrete gait time-series data are sample entropy (SE) and detrended fluctuation analysis (DFA), respectively [20, 21].

Both linear, as well as non-linear methods, generally require recording of a large number of consecutive strides (depending on the method between 50 – 500) in order to be measured reliably [20, 22-24]. In a confined laboratory space, consecutive strides can either be recorded using special overground (OG) protocols [22, 25] or on motorized treadmills. However, when using treadmills, it has been found that the constant speed (CS) of the treadmill imposes constraints to the gait and thereby reduces the magnitude of movement variability, as well as making the temporal structure of gait variability unrealistically regular as compared to OG walking [26-30]. It appears that the velocity constraint imposed by the treadmill impedes naturally occurring gait velocity fluctuations, which in return leads to the described distinct changes in magnitude and structure of gait variability. For example, in response to a given treadmill velocity, participants are required to modulate their step frequency or step length [31]. Therefore, a general recommendation is to avoid CS treadmills when observing gait variability outcomes.

As an alternative, self-paced (SP) treadmills have been developed, which continuously update the belt speed depending on the subject’s position on the treadmill and, thus, aim to match to the walking speed of the participant. This feedback based corrected belt speed, therefore, seeks to overcome the major limitation of traditional CS treadmills and can be implemented with varying degrees of sensitivity. For most non-variability gait outcomes, it has been found that SP and CS treadmill results are comparable, albeit some differences were found for linear variability measures [32, 33]. However, to the best of our knowledge, this is the first study investigating the validity of SP treadmills for the assessment of non-linear variability outcomes. It is conceivable that SP treadmills, due to their ability to allow for gait velocity fluctuations, allow for a more realistic, or OG-like, gait variability pattern to occur. Therefore, SP treadmills could potentially be a useful tool for a convenient data collection of long consecutive walking trials for the assessment of linear and non-linear gait variability outcomes.

Besides the limiting effect of CS treadmills on gait variability outcomes, it appears that their influence depends on the functioning of the motor control system and, therefore, might be exaggerated in patients with neuromotor deficits, for example, patients with Parkinson’s disease (PD). The typical shuffling and unsteady gait of PD corresponds to generally elevated levels of temporal and spatial magnitude of gait variability [34]. Furthermore, the temporal structure of gait variability is deteriorated in PD in that long-range correlation across consecutive gait cycles breaks down, and walking performance becomes more random and less structured [35-37]. One approach to manage Parkinsonian gait deficits is to provide PD with external rhythmic cues in order to re-gain a rhythmic pattern and, thus, normalization of the deteriorated gait variability [38-40]. Interestingly, CS treadmills have been used similarly in order to provide an external rhythmic cue and restore inadequate levels of gait variability in PD [41, 42]. Therefore, it is likely that the effect of CS treadmills on gait variability outcomes is larger in PD as compared to healthy control subjects (HC), as they provide additional external control for the malfunctioning nervous system to support walking function.

In light of the lack of information regarding the validity of SP treadmills to allow *natural* gait variability to occur, this study aimed to investigate (i) if spatio-temporal, linear and non-linear gait variability outcomes collected on SP treadmills are more similar to OG walking as compared to data collected on CS treadmills, (ii) which SP sensitivity setting might result in the most realistic gait variability patterns and (iii) if this effect is depending on the function of the nervous system, by comparing results for a group of HC as well as PD.

## Methods

### Participants

Thirteen HC subjects (6 females; mean [SD] age, height, and weight: 24.2 [3.9] years, 171.1 [8.5] cm, 71.5 [13.7] kg) and the same number of PD participated in the study. PD (8 females; mean [SD] age, height, and weight: 71.2 [7.3] years, 172.2 [11.4] cm, 70.4 [13.9] kg) were diagnosed on average for 6.9 [4.9] years and had a median [minimum; maximum] modified Hoehn & Yahr score of 2 [1; 2.5]. PD performed the experiment after intake of their regular anti-Parkinson medication (i.e., ON). All subjects provided written informed consent before participation, and the study protocol was approved by the Chapman University institutional review board and conducted in accordance with the declaration of Helsinki.

### Procedure

All participants performed at least a total of five walking trails, each of which was of 5-8 minutes, during one session. The first trial was an uninterrupted OG walking trial, outside along a 0.8km long, relatively straight pavement. The remaining four trials were randomized and performed on a self-paced treadmill (Motek Grail, Netherlands): CS walking (set at the pace determined during the initial OG trial), default SP mode (SP_default_), maximally low sensitive SP mode (SP_low_) and maximally high sensitive SP mode (SP_high_). The self-paced treadmill algorithm, a proportional-derivative controller, aims to keep a subject within the boundaries of a prescribed anterior-posterior 0.95m space around the center of the treadmill. In order to achieve this, the average trajectory position from four infra-red markers on the pelvis (i.e. right and left ASIS and PSIS, approximated CoM) is the central tenet in controlling the treadmill speed relative to the walking speed. The treadmill belt speed will be updated proportionally to a change in the subject’s position relative to the center of the treadmill and while considering the current gait speed. In general, an anterior deviation from the center will be responded with an increase in belt speed and a posterior position with a deceleration of the belt speed. The different sensitivity settings scale the response of the treadmill in terms of stronger (i.e. SP_high_) or weaker (i.e. SP_low_) response to a position change and more (i.e. SP_high_) or less (i.e. SP_low_) strong changes in belt speed [33, 43].

Before the treadmill trials, all participants got at least five minutes to familiarize themselves with walking in the self-paced mode. HC participants performed a second OG walking trial, which was randomized together with the treadmill trials, in order to estimate the intra-session reliability of the measurement system. During all trials, participants were instructed to maintain their preferred comfortable walking speed. Consecutive stride lengths (SL) and stride times (ST) from the dominant foot were recorded using a validated inertial-measurement unit system (APDM Mobility lab, USA) [44]. All data related to this study can be found in the supplementary material.

### Data analysis

For each trial, the first and last five steps were removed and all trials were cropped to the shortest trial of the subject in order to ensure the same data length in the subsequent analysis. The stride times and stride lengths time series data were then analyzed for linear gait variability magnitude, regularity and self-similarity by calculating the coefficient of variation (CV), SE and DFA, respectively. For SE we used a constant vector length (i.e., *m*) of 2 and similarity threshold (i.e., *r*) of 0.2 for all trials [45]. In order to investigate the sensitivity of the SE estimation to the input parameters, additional *m* and *r* combinations were tested (see supplementary material) [45]. For DFA, minimum and maximum window length was set at 16 and a ninth of the total data length, respectively [46].

### Statistical analysis

The results from the OG trials serve as the gold standard. All treadmill conditions are compared to the OG-trial, and the linear association between the two trials is assessed using the Pearson moment correlation coefficient (r). The method error (ME) provides a measure for the overall differences between the two trials in percent [47]. Whereas, the Bias and Limits of Agreement (LoA) from a Bland and Altman analysis represent systematic and random error effects between two trials, respectively [48]. A two-step approach will be used for determining the level of validity of each condition-outcome combination. First, only conditions with a significant correlation coefficient (based on the 95% confidence intervals) will be deemed acceptable. Second, the overall differences, systematic and random error will be considered.

## Results

### Benchmark intrasession reliability

HC performed on average 445 (±24) steps during the OG trials. The average OG gait speed of the group was 1.4 (±0.1) m/s. Except for DFA for SL, all conditions show significant correlations between the two OG trials. Intra-session reliability for stride time variability estimates was slightly better than for stride length, with a smaller amount of difference (ME) and systematic (Bias) and random error effects (LoA) (Table 1). SE showed the highest reliability of the three outcome measures.

**Table 1:** Intrasession reliability for the HC who performed two OG walking trials during the same day.

|  |  | r [95% CI] | ME [95% CI] | Bias (%) | LoA (%) |
| --- | --- | --- | --- | --- | --- |
| Stride time | CV | 0.79 [0.43; 0.94] | 9 [7; 16] | 0.0001 (<1) | 0.004 (26) |
|  | SE | 0.83 [0.53; 0.95] | 6 [4; 9] | 0.0109 (<1) | 0.221 (16) |
|  | DFA | 0.81 [0.47; 0.94] | 8 [6; 13] | -0.0139 (2) | 0.189 (22) |
| Stride length | CV | 0.63 [0.12; 0.88] | 10 [7; 16] | 0.0006 (3) | 0.005 (28) |
|  | SE | 0.69 [0.23; 0.90] | 4 [3; 7] | -0.0253 (1) | 0.235 (12) |
|  | DFA | 0.46 [-0.12; 0.81] | 10 [7; 17] | 0.0592 (6) | 0.263 (29) |

### Self-paced treadmill validation for HC

For ST-CV, only SP_default_ and SP_low_ result in significant correlations. All conditions for ST-SE resulted in significant correlations. Here, the SP_default_ resulted in the least overall, systematic and random error of 8, 1, and 21%, respectively. None of the ST-DFA outcomes was significantly correlated (Table 2). For SL no condition resulted in a significant correlation between OG and treadmill trials.

**Table 2:** Statistical comparison of overground and treadmill trials for the HC group. Temporal and spatial gait outcomes and three variability measures. Bold: Results with significant correlation coefficients.

|  |  |  | r [95% CI] | ME [95% CI] | Bias (%) | LoA (%) |
| --- | --- | --- | --- | --- | --- | --- |
| Stride time | CV | OG – CS | 0.23 [-0.37; 0.69] | 18 [13; 30] | 0.001 (8) | 0.007 (50) |
|  |  | OG – SP<sub>default</sub> | **0.71 [0.25; 0.91]** | **17 [13; 29]** | **-0.002 (10)** | **0.008 (49)** |
|  |  | OG – SP<sub>high</sub> | 0.49 [-0.08; 0.82] | 16 [11; 26] | -0.002 (9) | 0.007 (43) |
|  |  | OG – SP<sub>low</sub> | **0.60 [0.07; 0.86]** | **14 [11; 24]** | **-0.001 (4)** | **0.006 (41)** |
|  | SE | OG – CS | **0.62 [0.11; 0.87]** | **8 [6; 14]** | **-0.005 (3)** | **0.344 (24)** |
|  |  | OG – SP<sub>default</sub> | **0.74 [0.31; 0.92]** | **8 [5; 12]** | **0.010 (1)** | **0.297 (21)** |
|  |  | OG – SP<sub>high</sub> | **0.66 [0.17; 0.89]** | **8 [6; 13]** | **-0.042 (3)** | **0.321 (22)** |
|  |  | OG – SP<sub>low</sub> | **0.57 [0.03; 0.85]** | **10 [7; 16]** | **0.039 (3)** | **0.381 (27)** |
|  | DFA | OG – CS | 0.50 [-0.07; 0.83] | 12 [9; 21] | 0.102 (13) | 0.281 (35) |
|  |  | OG – SP<sub>default</sub> | -0.07 [-0.60; 0.50] | 18 [13; 30] | -0.051 (6) | 0.452 (51) |
|  |  | OG – SP<sub>high</sub> | 0.44 [-0.14; 0.80] | 13 [9; 21] | 0.057 (7) | 0.294 (35) |
|  |  | OG – SP<sub>low</sub> | 0.13 [-0.45; 0.63] | 16 [11; 26] | -0.075 (8) | 0.392 (44) |
| Stride length | CV | OG – CS | -0.28 [-0.72; 0.32] | 24 [17; 39] | 0.003 (20) | 0.012 (66) |
|  |  | OG – SP<sub>default</sub> | 0.31 [-0.29; 0.74] | 22 [15; 36] | -0.005 (23) | 0.013 (60) |
|  |  | OG – SP<sub>high</sub> | 0.46 [-0.13; 0.81] | 13 [10; 22] | -0.007 (29) | 0.009 (37) |
|  |  | OG – SP<sub>low</sub> | 0.27 [-0.33; 0.72] | 16 [12; 27] | -0.003 (15) | 0.010 (45) |
|  | SE | OG – CS | 0.37 [-0.23; 0.76] | 7 [5; 11] | 0.045 (2) | 0.364 (19) |
|  |  | OG – SP<sub>default</sub> | -0.05 [-0.58; 0.52] | 12 [9; 20] | 0.057 (3) | 0.647 (34) |
|  |  | OG – SP<sub>high</sub> | 0.49 [-0.08; 0.82] | 6 [4; 9] | -0.150 (7) | 0.310 (15) |
|  |  | OG – SP<sub>low</sub> | 0.38 [-0.22; 0.77] | 7 [5; 12] | -0.028 (1) | 0.390 (20) |
|  | DFA | OG – CS | -0.43 [-0.80; 0.15] | 20 [14; 33] | 0.260 (32) | 0.448 (55) |
|  |  | OG – SP<sub>default</sub> | 0.22 [-0.37; 0.69] | 14 [10; 24] | -0.045 (5) | 0.387 (40) |
|  |  | OG – SP<sub>high</sub> | 0.15 [-0.44; 0.65] | 18 [13; 29] | 0.196 (23) | 0.411 (49) |
|  |  | OG – SP<sub>low</sub> | -0.12 [-0.63 ; 0.46] | 15 [11; 24] | -0.248 (23) | 0.436 (41) |

### Self-paced treadmill validation for PD

The PD performed significantly fewer steps during the same trial time; on average 263 (±87) steps. The average OG gait speed of the PD group was 1.2 (±0.2) m/s. For ST-CV only the SP_low_ condition resulted in a significant correlation to OG-trials. None of the ST-SE trials was significantly correlated. SP_default_ revealed a significant correlation for ST-DFA, with ME, Bias and LoA of 33, 31 and 90%, respectively (Table 3).

**Table 3:** Statistical comparison of overground and treadmill trials for the PD group. Temporal and spatial gait outcomes and three variability measures. Bold: Results with significant correlation coefficients.

|  |  |  | r [95% CI] | ME [95% CI] | Bias (%) | LoA (%) |
| --- | --- | --- | --- | --- | --- | --- |
| Stride time | CV | OG – CS | 0.16 [-0.43; 0.65] | 56 [40; 92] | -0.008 (35) | 0.040 (155) |
|  |  | OG – SP<sub>default</sub> | -0.15 [-0.65; 0.44] | 40 [29; 67] | -0.009 (37) | 0.029 (112) |
|  |  | OG – SP<sub>high</sub> | 0.20 [-0.39; 0.67] | 36 [26; 60] | -0.014 (50) | 0.029 (101) |
|  |  | **OG – SP<sub>low</sub>** | **0.73 [0.27; 0.92]** | **40 [28; 68]** | **-0.011 (42)** | **0.029 (111)** |
|  | SE | OG – CS | 0.13 [-0.45; 0.64] | 17 [12; 28] | -0.000 (<1) | 0.829 (47) |
|  |  | OG – SP<sub>default</sub> | 0.36 [-0.24; 0.76] | 16 [11; 26] | 0.115 (7) | 0.747 (44) |
|  |  | OG – SP<sub>high</sub> | 0.53 [-0.03; 0.83] | 16 [12; 27] | 0.219 (13) | 0.747 (45) |
|  |  | OG – SP<sub>low</sub> | 0.32 [-0.31; 0.76] | 18 [13; 30] | 0.146 (9) | 0.822 (49) |
|  | DFA | OG – CS | -0.10 [-0.62; 0.47] | 142 [101; 234] | 0.468 (60) | 3.074 (394) |
|  |  | **OG – SP<sub>default</sub>** | **0.64 [0.14; 0.88]** | **33 [23; 54]** | **-0.378 (31)** | **1.085 (90)** |
|  |  | OG – SP<sub>high</sub> | 0.28 [-0.32; 0.72] | 34 [24; 55] | -0.084 (8) | 0.982 (93) |
|  |  | OG – SP<sub>low</sub> | -0.58 [-0.87; -0.01] | 78 [55; 132] | 0.389 (47) | 1.785 (215) |
| Stride length | CV | OG – CS | **0.62 [0.11; 0.87]** | **39 [28; 64]** | **-0.004 (12)** | **0.031 (108)** |
|  |  | OG – SP<sub>default</sub> | 0.30 [-0.30; 0.73] | 34 [24; 56] | -0.021 (56) | 0.036 (95) |
|  |  | OG – SP<sub>high</sub> | 0.07 [-0.50; 0.59] | 78 [56; 128] | -0.042 (87) | 0.103 (215) |
|  |  | **OG – SP<sub>low</sub>** | **0.70 [0.20; 0.91]** | **35 [24; 59]** | **-0.020 (55)** | **0.035 (96)** |
|  | SE | OG – CS | **0.76 [0.36; 0.92]** | **14 [10; 24]** | **-0.141 (7)** | **0.808 (40)** |
|  |  | OG – SP<sub>default</sub> | 0.40 [-0.20; 0.78] | 20 [14; 33] | 0.470 (27) | 0.950 (55) |
|  |  | OG – SP<sub>high</sub> | -0.20 [-0.68; 0.39] | 33 [24; 54] | 0.631 (38) | 1.509 (91) |
|  |  | **OG – SP<sub>low</sub>** | **0.63 [0.09; 0.89]** | **21 [15; 36]** | **0.266 (15)** | **1.057 (59)** |
|  | DFA | OG – CS | -0.54 [-0.84; 0.02] | 83 [59; 137] | 0.240 (30) | 1.835 (229) |
|  |  | OG – SP<sub>default</sub> | 0.35 [-0.25; 0.75] | 29 [21; 48] | -0.588 (48) | 0.971 (80) |
|  |  | OG – SP<sub>high</sub> | 0.32 [-0.28; 0.74] | 20 [15; 33] | -0.512 (44) | 0.660 (56) |
|  |  | OG – SP<sub>low</sub> | 0.52 [-0.08; 0.84] | 31 [22; 53] | -0.247 (24) | 0.904 (86) |

For SL-CV only CS and SP_low_ revealed a significant correlation, with CS having a lower systematic error of 12%, as compared to 55% in SP_low_ (Figure 1). Similarly, for SL-SE only CS and SP_low_ result in significantly correlated results, with CS showing lower ME, Bias and LoA of 14, 7, 40%, respectively. None of the SL-DFA conditions resulted in significant correlations.

**Figure 1:**
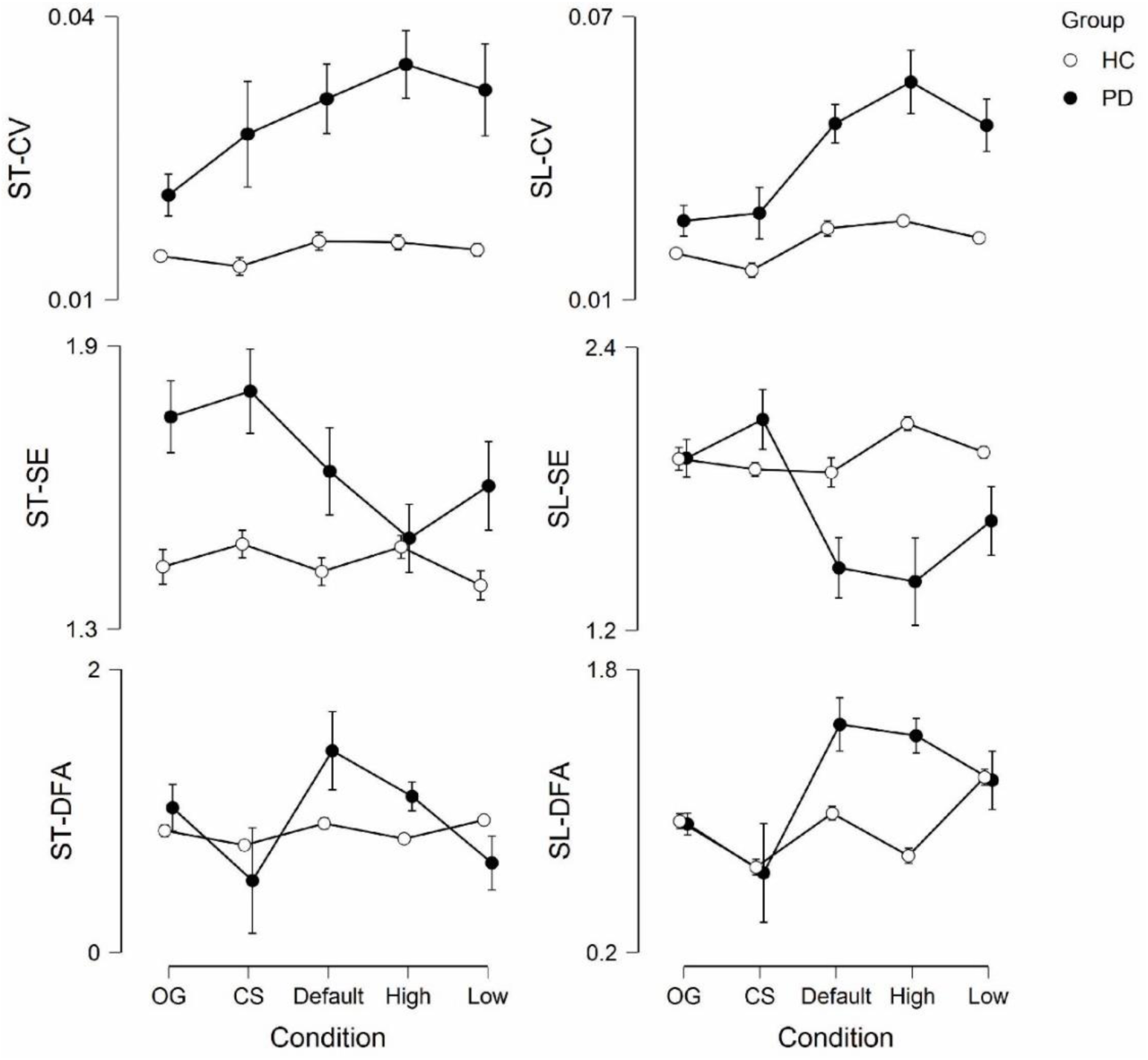
Results for three variability measures across five walking conditions for healthy control subjects and Patients with Parkinson’s disease.

## Discussion

This study investigated if self-paced treadmills avoid the limitations imposed by constant speed treadmills during the assessment of linear and non-linear gait variability outcomes. We find that for healthy young subjects, stride time outcomes are most similar to overground walking in the default self-paced mode for the coefficient of variation and sample entropy. Detrended fluctuation analysis results in invalid observations. For stride length outcomes, none of the outcome-condition combinations resulted in valid observations. For the individuals with Parkinson’s disease, coefficient of variation, and detrended fluctuation analysis of stride time measures are most similar to overground in the low sensitivity or default sensitivity mode. The regularity of stride times, as measured by sample entropy, cannot be measured validly in any treadmill condition in this cohort. Stride length outcomes, however, are most similar to overground walking in the constant speed condition.

The validity of gait variability outcomes derived from treadmill walking also depends on the general reliability of the assessment of the outcome under observation. It is known that a larger number of steps is required for the reliable evaluation of gait variability as compared to mean measures of gait. For linear measures (e.g. coefficient of variation), about 50 steps have been found to result in reliable variability estimates for asymptomatic persons and PD [22, 25]. For sample entropy and DFA, on the other hand, it was suggested to include not less than 200 and 500 samples for reliable estimates, respectively [20, 24]. Therefore, estimates for CV and SE should be considered reliable in both groups. However, the poor validity of DFA results might be the result of too short time-series used in the analysis.

The regulatory effect of motorized treadmills, that act as an external pacemaker, on patients with Parkinson’s disease has been documented before. Similarly to Warlop and colleagues, we find markedly elevated levels of temporal magnitude of variability in PD during treadmill walking that is absent in healthy controls [49]. Also, the generally poorer validity in the PD for all outcomes indicates that treadmill use is particularly problematic for the assessment of patient cohorts with sensory-motor deficits. For example, the average effect size (Cohen’s d) from comparing stride time variability outcomes in PD and HC is about 0.63 and the effect of antiparkinson medication on temporal gait variability measures was found to be 0.66 [34, 50]. In this study, the effect of the treadmill on the CV of stride time of PD in the best condition (i.e., SP_low_) was 0.78. For the overall most similar condition (i.e., spatial sample entropy during CS treadmill walking), the effect size is still 0.3, or about 50% of the disease or medication effect. Thus, using any motorized treadmill for the assessment of gait variability in PD is not recommended.

In conclusion, for healthy subjects, the default self-paced mode increases the validity of temporal gait variability outcomes (except for DFA results), as compared to constant speed treadmills. But spatial variability outcomes cannot be measured reliably. For patients with Parkinson’s disease, temporal gait variability outcomes are best assessed during the low and default self-paced mode, albeit with generally substantial differences to overground walking of 30-40%. Spatial gait variability in patients is best assessed using the constant speed mode. However, the effects of even the best treadmill setting are so large that they potentially mask the effects induced by interventions to these patients and, therefore, using constant or self-paced treadmills for the assessment of gait variability outcomes in these patients cannot be recommended.

## Supporting information

Sample Entropy calculation

## Acknowledgements

The authors would like to thank the study participants for their time and effort, as well as Kaitlyn Raymundo and Courtney Justus for their assistance during the conduct of the measurements.

## Supplementary material

All stride time and stride length data used in this study, as well as the comparison of different input parameter choices during the SE analysis can be found at: https://doi.org/10.36837/chapman.000126

