## Supplementary material for "Self-paced treadmills do not allow for valid observation of linear and non-linear gait variability outcomes in patients with Parkinson’s disease": Sample Entropy calculation

**Comparison of input parameters for the sample entropy calculation:**

The stride time and stride length data from the healthy control participants during overground (OG) and constant speed treadmill (CS) walking was used to investigate the sensitivity of sample entropy (SE) calculations to different *m* and *r* combinations. The vector length *m* was either 2 or 3, and the similarity threshold *r* was set at 0.15, 0.2, 0.25, or 0.3.

*Stride time*

As can be seen in Figure 1, the choice of *m* and *r* did not influence the resulting SE, in particular when comparing OG and CS walking in terms of their regularity.

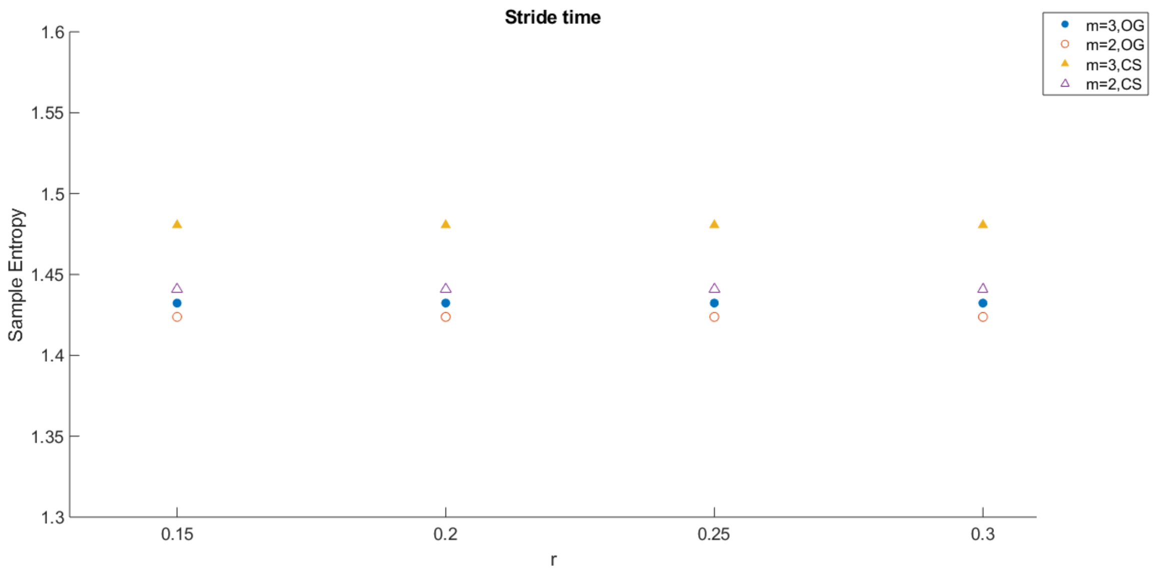


*Figure 1: Results for sample entropy as a function of varying m and r combinations from stride time data.*

*Stride length*

For stride length outcomes, Figure 2 shows that results are affected by the *m* and *r* choice. OG walking was least regular for *m* = 2 and *r* = 0.15, 0.2 and 0.25. When choosing *m* = 2 (or 3) and *r* = 0.3, however, OG becomes more regular as compared to CS walking. The fact that SE can be sensitive to the choice of *r* has been reported before [1]. As the most consistent results are found for *r*-values below 0.3, we decided to apply *m* = 2 and *r* = 0.2 for all SE calculations.

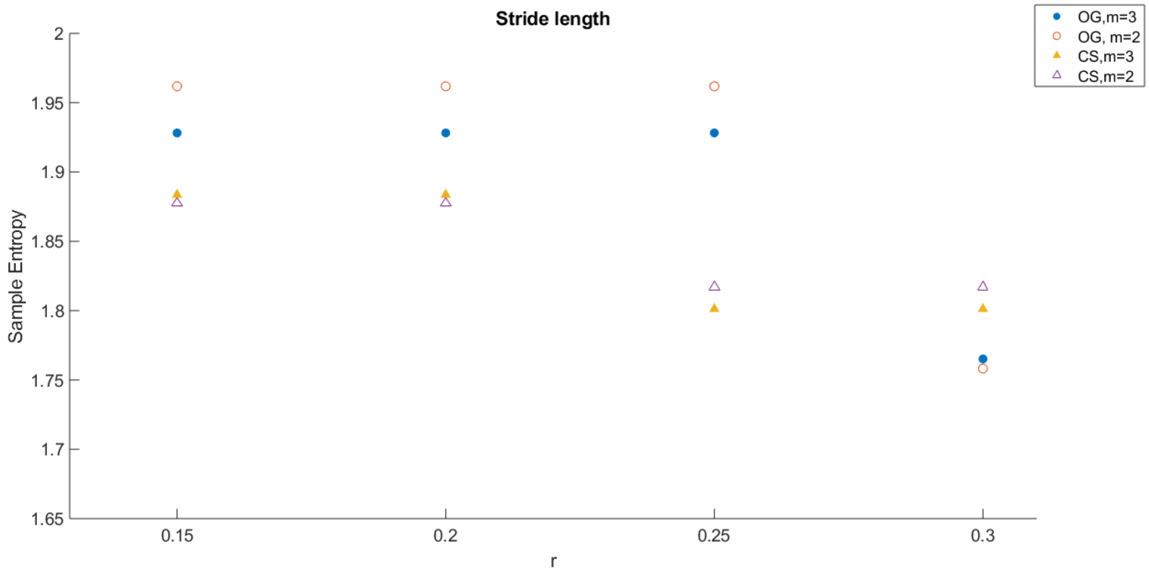


*Figure 2: Results for sample entropy as a function of varying m and r combinations from stride length data.*
